# Selection of appropriate metagenome taxonomic classifiers for ancient microbiome research

**DOI:** 10.1101/260042

**Authors:** Irina M. Velsko, Laurent A. F. Frantz, Alexander Herbig, Greger Larson, Christina Warinner

**Affiliations:** The Palaeogenomics and Bio-Archaeology Research Network, Research Laboratory for Archaeology and the History of Art, University of Oxford, Oxford, UK; School of Biological and Chemical Sciences, Queen Mary University of London, London, UK; Department of Archaeogenetics, Max Planck Institute for the Science of Human History, Jena, Germany; Department of Anthropology, University of Oklahoma, Norman, Oklahoma, USA; Department of Periodontics, University of Oklahoma Health Sciences Center, Oklahoma City, Oklahoma, USA

**Author notes:** Corresponding Author: Christina Warinner, Max Planck Institute for the Science of Human History, Kahlaische Strasse 10, Jena, Germany 07745. Current affiliation: Department of Biological Sciences, Clemson University, Clemson, SC, USA.

## Abstract

Metagenomics enables the study of complex microbial communities from myriad sources, including the remains of oral and gut microbiota preserved in archaeological dental calculus and paleofeces, respectively. While accurate taxonomic assignment is essential to this process, DNA damage, characteristic to ancient samples (*e.g.* reduction in fragment size), may reduce the accuracy of read taxonomic assignment. Using a set of *in silico*-generated metagenomic datasets we investigated how the addition of ancient DNA (aDNA) damage patterns influences microbial taxonomic assignment by five widely-used profilers: QIIME/UCLUST, MetaPhlAn2, MIDAS, CLARK-S, and MALT (BLAST-X-mode). *In silico*-generated datasets were designed to mimic dental plaque, consisting of 40, 100, and 200 microbial species/strains, both with and without simulated aDNA damage patterns. Following taxonomic assignment, the profiles were evaluated for species presence/absence, relative abundance, alpha-diversity, beta-diversity, and specific taxonomic assignment biases. Unifrac metrics indicated that both MIDAS and MetaPhlAn2 provided the most accurate community structure reconstruction. QIIME/UCLUST, CLARK-S, and MALT had the highest number of inaccurate taxonomic assignments; however, filtering out species present at <0.1% abundance greatly increased the accuracy of CLARK-S and MALT. All programs except CLARK-S failed to detect some species from the input file that were in their databases. Ancient DNA damage resulted in minimal differences in species detection and relative abundance between simulated ancient and modern datasets for most programs. In conclusion, taxonomic profiling biases are program-specific rather than damage-dependent, and the choice of taxonomic classification program to use should be tailored to the research question.

**Importance:** Ancient biomolecules from oral and gut microbiome samples have been shown to preserve in the archaeological record. Studying ancient microbiome communities using metagenomic techniques offer a unique opportunity to reconstruct the evolutionary trajectories of microbial communities through time. DNA accumulates specific damage over time, which could potentially affect taxonomic classification and our ability to reconstruct community assemblages accurately. It is therefore necessary to assess whether ancient DNA (aDNA) damage patterns affect metagenomic taxonomic profiling. Here, we assessed biases in community structure, diversity, species detection, and relative abundance estimates by five popular metagenomic taxonomic classification programs using *in silico*-generated datasets with aDNA damage. Age-related damage patterns had minimal impact on the taxonomic profiles produced by each program, and biases were intrinsic to each program. Therefore, an appropriate classification program should be chosen that minimizes the biases related to the questions being addressed.

## Introduction

Ancient microbiome research offers the possibility of tracing the evolution of the complex microbial communities that play an integral role in shaping population health and disease. Palaeomicrobiology uses archaeological material to trace the emergence and spread of microorganisms throughout history and prehistory. Archaeological dental calculus and palaeofeces are promising substrates for ancient human microbiome studies, as they have been shown to preserve DNA (1), proteins (1, 2), and small molecule metabolites (3) from the resident microbes and the host. During life, these dense microbial communities contain hundreds of species, predominantly composed of bacteria (4), but also including archaea (4), viruses (5), fungi (6), and protists (7). Characterizing the microbial ecology of host-associated microbiota through time is a necessary step in understanding the function of these microbial communities, and further how they interact with the host.

DNA in archaeological samples, including ancient microbial samples, acquires predictable age-related damage patterns, including short fragment lengths (typically <100 bp) (8) with break-points coinciding with depurination, and accumulation of cytosine to thymine transitions at the ends of the molecules (8). The ubiquity and predictability of these damage patterns means that they are often used to authenticate ancient DNA and estimate modern DNA contamination (9, 10), and the short fragment lengths of ancient DNA negate the need for shearing during library construction for high throughput sequencing (HTS). These same properties, however, potentially affect taxonomic classification of microbial DNA sequence reads more difficult, or less accurate. Reads that are too short, for example, may not be specific enough for classification at the taxonomic level desired. Cytosine to thymine transitions may also cause misclassification or prevent classification, such that reads may be misleadingly assigned to unidentified taxa, thereby inflating diversity estimates. Additionally, although 16S rRNA gene amplicon sequencing is popular for profiling complex microbial communities, taxon-specific length polymorphisms in this gene combined with the relatively long lengths of the hypervariable regions (>150 bp), make it problematic for sequencing degraded DNA from ancient microbial communities (8). Instead, shotgun metagenomic sequencing, which is highly compatible with short DNA fragments, is the preferred analytical approach for ancient microbiome samples (1, 11).

Community profiling by DNA shotgun sequencing is currently the most comprehensive method used to assess microbiome community composition, and a variety of computational tools are available to reconstruct the species present from the millions of short sequences that comprise HTS datasets. There are several methods for taxonomic assignment available. Popular methods include matching reads to 16S rRNA gene sequences (QIIME (12), Mothur (13), or to single-copy gene panels (MetaPhlAn2 (14, 15), MIDAS (16), PhyloSift (17)), k-mer-based whole-genome matching (Kraken (18), CLARK (19, 20), and hybrid k-mer-based matching and alignment extension (MALT (21, 22)). While there are several publications comparing the accuracy, specificity, and precision of various metagenomic classification programs for modern samples (e.g., (23–25)), no study has yet compared the performance of these approaches on ancient DNA.

In order to assess the performance of metagenomic classification systems on ancient DNA, we performed a comparison of the community profile of six metagenomic classification programs that use different taxonomic assignment methods (QIIME, DADA2 (26), MetaPhlAn2, MIDAS, CLARK-S and MALT in BLAST-X-mode). We used *in silico*-generated ancient and modern metagenome samples to estimate the accuracy of these programs. Our results indicate that the effect of DNA damage patterns on taxonomic assignments is variable across programs. We show, however, that most of the programs tested here are robust to misassignment due to DNA damage. Overall, our results indicate that taxonomic assignment biases are similar between modern and ancient simulated metagenomic samples.

## Results

### Description of the datasets

A total of 39 *in silico* generated metagenomic community samples were generated by independent runs of gargammel (27) (Supplemental Table S1). Three overlapping sets of genomes were used as input: one set had 40 genomes, the second had 100 genomes, and the third had 200 genomes. All genomes in the 40 genome set were included in the 100 genome set, and all genomes in the 100 set were included in the 200 set. Each genome was represented in equal abundance, where in the 40, 100, and 200 genome datasets each genome comprises 2.5%, 1%, and 0.5% of the total DNA, respectively. There were 13 independent samples for each set of genomes, where ten replicates had simulated aDNA damage patterns (ancient dataset) and three replicates did not have aDNA damage patterns (modern dataset). The estimated copy number of each genome in each dataset is presented in Supplemental Table S1. We additionally filtered the output profiles to remove species present at <0.1% abundance to understand how filtering low-abundance, often false-positive, taxa affected diversity metrics. The cut-off of 0.1% was arbitrarily selected based on (23).

### Community structure is consistent between ancient and modern simulated datasets

We first sought to determine if any of the taxonomic classification programs produced a community structure that closely resembled the true input files by measuring beta-diversity. We used both weighted UniFrac (phylogenetic relatedness accounting for relative abundance of organisms) and unweighted UniFrac metrics (phylogenetic relatedness without accounting for relative abundance of organisms) on full and filtered (>0.1% abundance) tables. Principal coordinates analysis (PCoA) of the beta-diversity metrics were plotted to visualize relatedness between community structure of the input files and community structure as determined by each of the 5 programs tested (Fig. 1A, B), and demonstrated that classification of replicate samples was highly consistent by each program, although QIIME showed the greatest variance between replicates. Filtering low-abundance species did not affect the weighted UniFrac distance, as this metric accounts for relative abundance of species, and therefore removing low-abundance species minimally affects the final score. Additionally, there was very little difference in the scores of the ancient and modern datasets for all programs, although QIIME/UCLUST demonstrated the greatest age-related difference in beta-diversity. MIDAS-determined community structure calculated by weighted UniFrac distance was most similar to the input files for 40 and 100-genome datasets (Fig. 1A). CLARK-S and MALT community structures were more similar to each other than to any of the other programs for all datasets, while the community structures reconstructed using QIIME/UCLUST and MetaPhlAn2 were each distinct from the other programs and did not plot near any other programs in the PCoA (Fig. 1A). Using the non-phylogenetic abundance-weighted Bray-Curtis distance we observed similar PCoA plotting patterns by each group, relative to the true input, at the species and genus levels (Figs. S2-S4).

**Figure 1.**
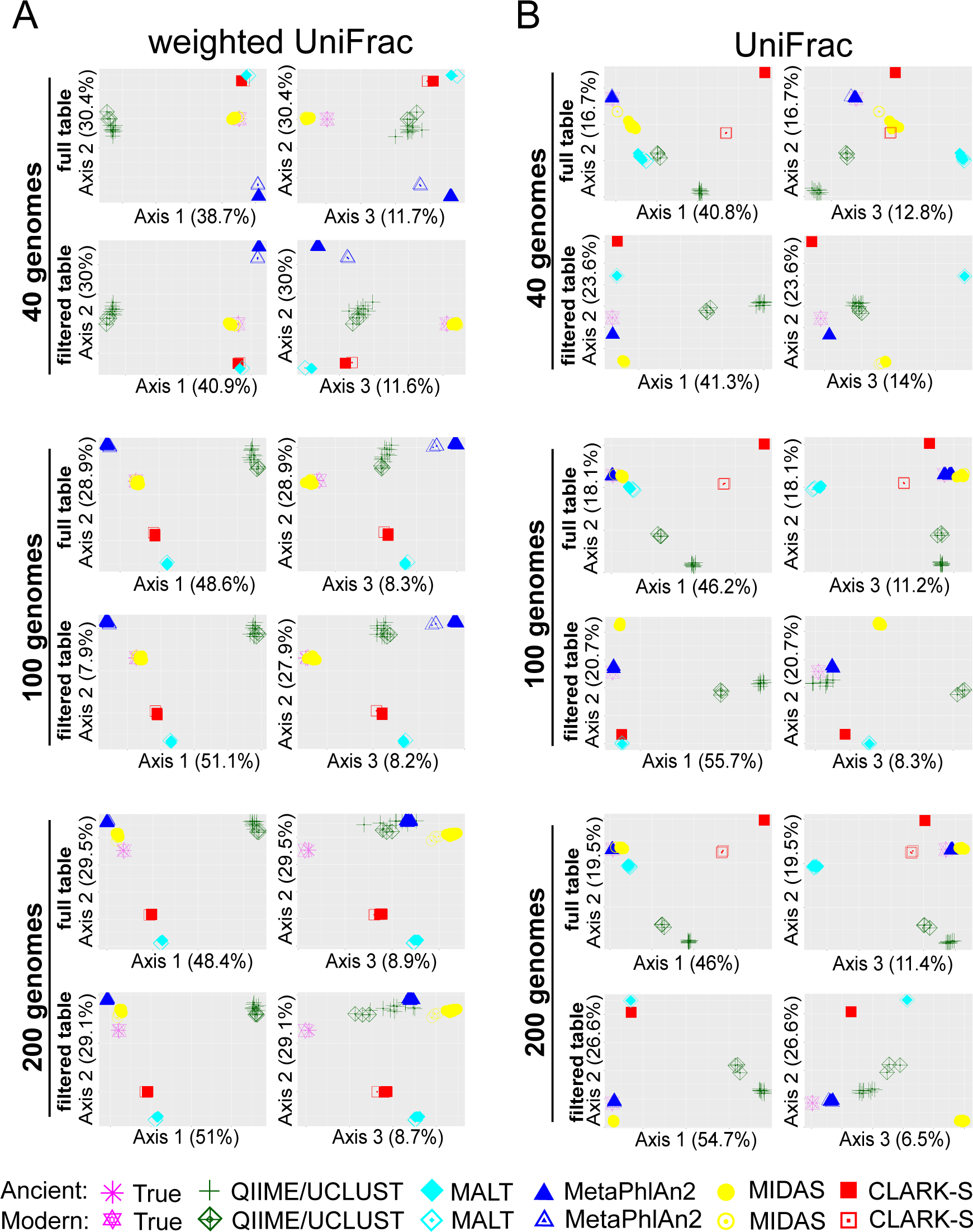
Age-related damage patterns minimally influence reported phylogenetic-based community structure. (A) Principal coordinates analysis plots of abundance-weighted UniFrac beta-diversity for datasets made with 40, 100, and 200 genomes for full output tables and tables filtered to remove species present at < 0.1% abundance. (B) Principal coordinates analysis plots of UniFrac beta-diversity for datasets made with 40, 100, and 200 genomes for full output tables and tables filtered to remove species present at < 0.1% abundance.

**Figure 2.**
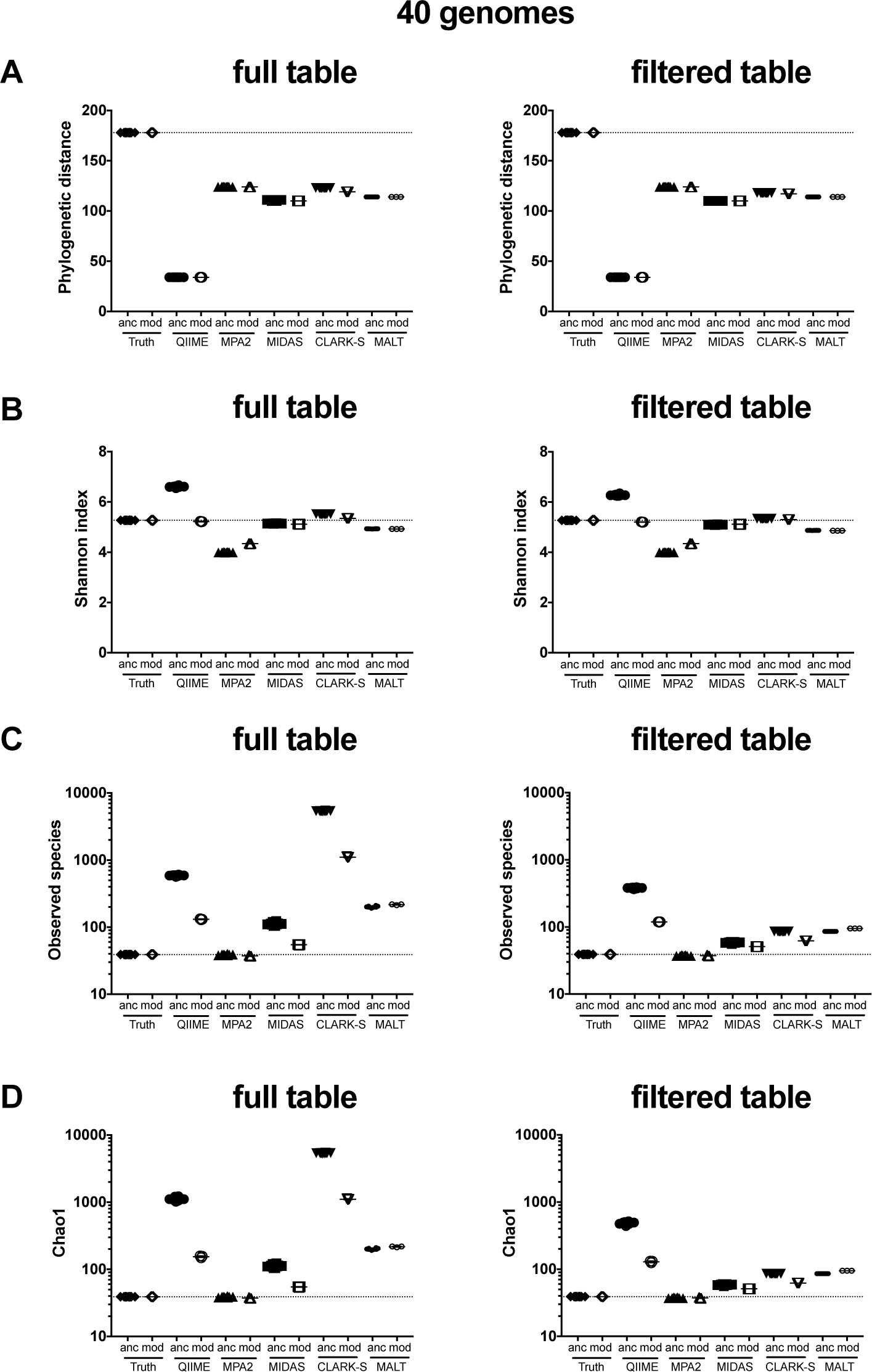
Age-related damage patterns slightly increase within-sample diversity. Alpha diversity of 40-genome datasets calculated by (A) Faith’s phylogenetic distance, (B) Shannon index, (C) Observed species, and (D) Chao1 for full output tables and tables filtered to remove species present at < 0.1% abundance. MPA2 - MetaPhlAn2, anc - ancient simulated dataset, mod - modern simulated dataset.

Plots of beta-diversity by the standard (unweighted) UniFrac metric, which accounts for species presence/absence but not abundance, were distinct from the weighted UniFrac plots, demonstrating differences in the ability of the five programs to accurately reflect the species composition *vs* composition plus abundance (Fig. 1B). Filtering out species present at <0.1% abundance noticeably altered the relationship of the programs to each other in the PCoA plots.

CLARK-S and QIIME/UCLUST exhibited substantial differences in community structure between ancient and modern datasets. Filtering removed this difference only for CLARK-S, while QIIME/UCLUST modern and ancient datasets remained distinctly plotted, suggesting that QIIME/UCLUST reported several taxa not in the input files at higher abundance than the cut-off of 0.1%. In contrast to the weighted UniFrac PCoA plots, MetaPhlAn2 community structure was most similar to truth for 40-, 100-, and 200-genome datasets, filtered and full tables, followed by MIDAS. Filtering output tables reduced the community structure similarity between MIDAS and the true input, and makes the community structure of CLARK-S and MALT more similar to each other, suggesting the most abundant species are detected in similar proportions by CLARK-S and MALT. Using the non-phylogenetic Jaccard distance we observed similar PCoA plotting patterns by each program, relative to the true input, at the species and genus levels (Figs. S2-S4).

### Community diversity is program-dependent

To understand the differences in community structure we observed in beta-diversity analyses, we assessed the alpha-diversity of the communities produced by the five taxonomic classification programs, using several metrics to account for different components of community diversity. Faith’s phylogenetic distance (PD), which determines the community diversity based on the phylogenetic relatedness of the species present, was estimated to be much lower than the true PD by all of the programs for the 40-, 100-, and 200-genome datasets, full and filtered tables, ancient and modern simulations (Figs. 2A, S5A, S6A). QIIME/UCLUST generated the lowest PD, while MetaPhlAn2 and CLARK-S were both slightly higher than MIDAS and MALT. CLARK-S was the only program with a slight difference in PD between ancient and modern simulated datasets, but when the table was filtered the modern and ancient sample diversity was equivalent.

The Shannon index, which accounts for species presence/absence and evenness, showed little difference between ancient and modern simulated datasets per program and was unaffected by filtering (Figs. 2B, S5B, S6B). As the number of genomes in the input files (truth) increased, the Shannon index values for each program decrease relative to true value (*i.e*., in the 40-genome set QIIME/UCLUST, MIDAS, and CLARK-S are above true value, and in the 200-genome set all program values are below the true value). This may be caused by the fact that the Shannon index of communities with dominant species is expected to be lower than those with even abundance across species, even if the former communities is more species rich.

The observed species is the total number of species/subspecies detected by each program (except QIIME/UCLUST which included all OTUs because it poorly resolves species-level differences). QIIME/UCLUST, MIDAS, CLARK-S, and MALT always overestimate the total number of species in the samples (by 1X-150X), and the number of estimated species/subspecies in ancient simulated samples is much higher than in modern simulated samples for QIIME/UCLUST and CLARK-S, and to a lesser extent MIDAS (Figs. 2C, S5C, S6C). Filtering reduced the number of observed species by CLARK-S substantially, by MALT and MIDAS slightly, and by QIIME/UCLUST minimally. In contrast to the other programs, MetaPhlAn2 slightly underestimates the total number of species in all of the datasets, and is consistently closest to the true number. Chao1 diversity metrics, which include an estimation of undetected species in the sample, exhibited very similar patterns to observed species for all programs (Figs. 2D, S5D, S6D).

### Individual program performance and biases

We next assessed how well each program detected the presence and abundance of species present in both modern and ancient simulated datasets. To do so, we calculated the true and inferred relative abundance of each input genome for each of the five programs, and determined the percent over- or under- assignment (Fig. 3, S7-S8). Given the limited species-level resolution afforded by QIIME/UCLUST, we limited our analysis to genus-level assignments for this program. MetaPhlAn2 is does not distinguish between several species *(i.e. Streptococcus mitis* and *S. oralis)* because their marker genes are indistinguishable, and the relative abundance of these in the input files was likewise combined for calculations. Generally, the species detected/not detected are consistent between ancient/modern simulated datasets, as is the percent and direction of over/under-estimation. We have additionally presented as bar charts (Fig. 4, S9-S18) the relative abundance of each species in the input (labeled If, “Input fastq/a”, and 16f “Input 16S rRNA gene-identified read fastq/a”) and output profiles from each program. The first output profile bar (labeled Id, “Input species detected”) excludes the false-positive species not in the input files (grouped together as “other” assignments). The second output profile bar (labeled Ad, “All species detected”) includes the “other” assignments to visualize how skewed the proportions of input species are by assignments to taxa not in the input. Assessment of each of the programs is included in the program-specific sections below.

**Figure 3.**
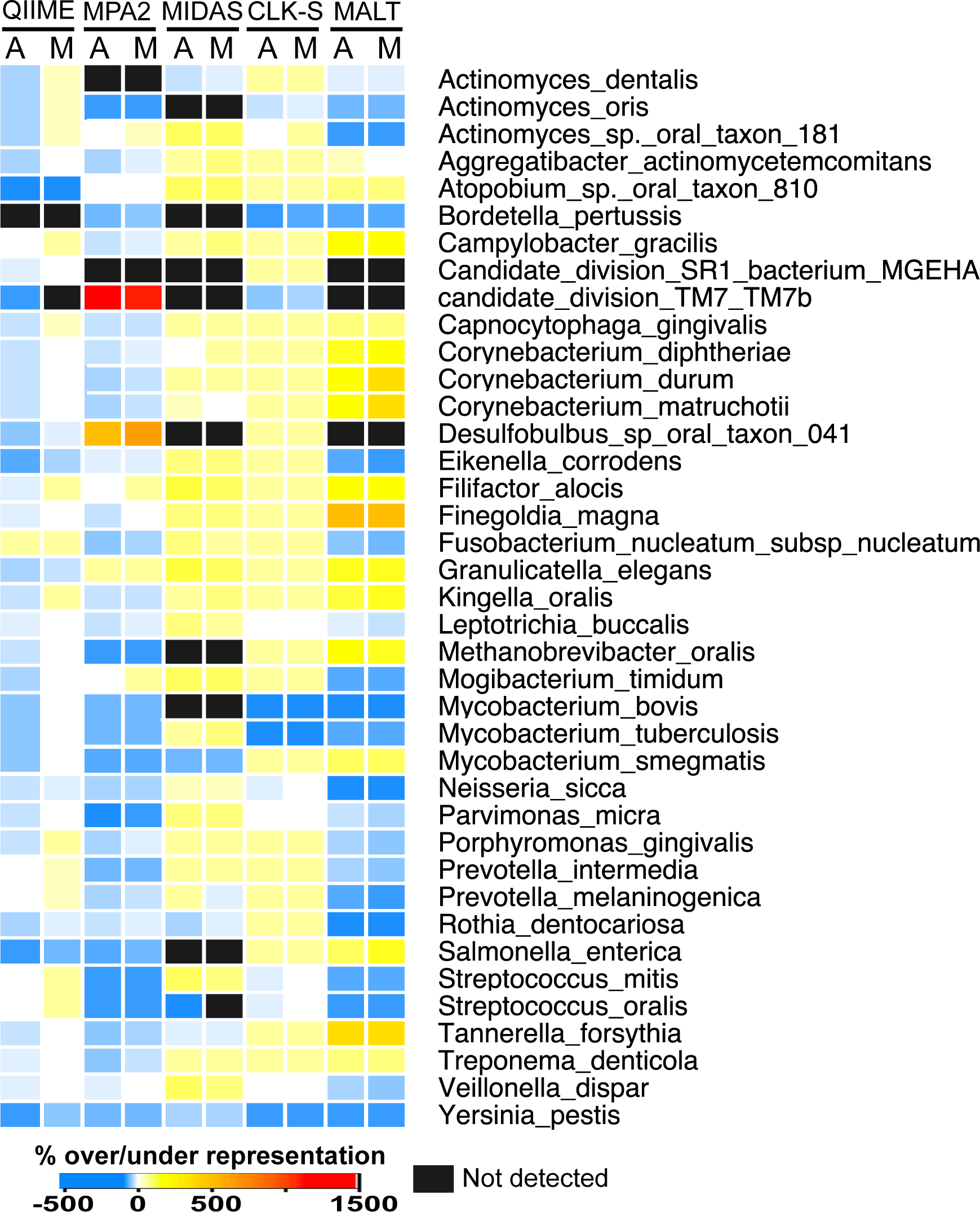
Species detection and over/under-representation differ by program but not age-related damage. Heat-map showing for each program tested the species relative abundance under-represented (blues), over-represented (yellows, oranges, reds), not detected (black), and accurately represented (white) relative to the true input files for modern and ancient 40-genome datasets. Where programs were unable to distinguish species, strains, or subspecies a single bar across those genomes is colored to represent the over/under-representation of the lowest identifiable taxonomic level. MPA2 - MetaPhlAn2, CLK-S - CLARK-S; A - ancient simulated dataset, M - modern simulated dataset.

**Figure 4.**
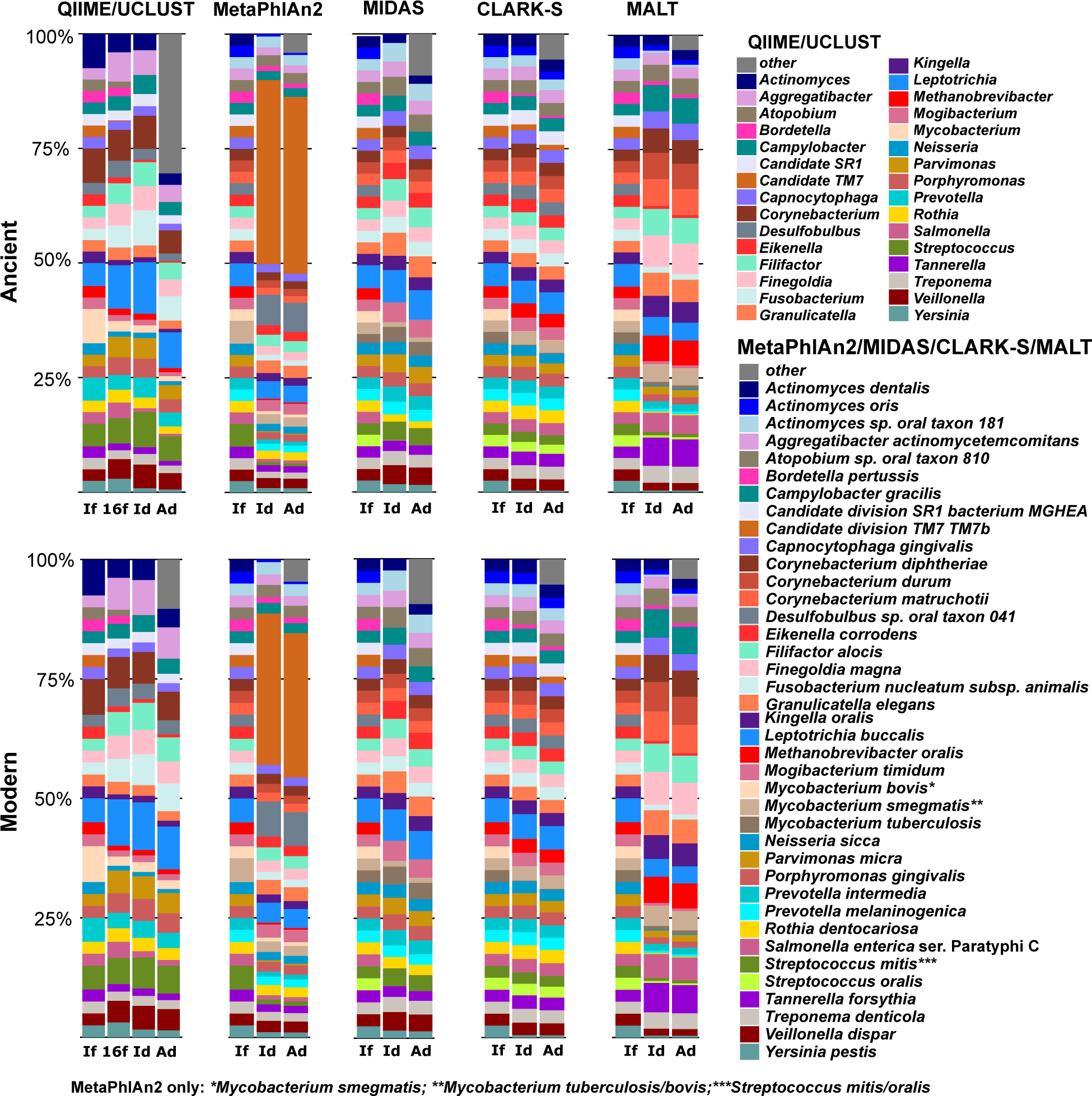
Differences in species relative abundance are program-specific and minimally affected by age-related damage. Program-specific differences in species detection and relative abundance are consistent between ancient (top) and modern (bottom) 40-genome simulated datasets. Relative abundances of each bar represent: If - true input fasta file, Id - input species detected, and Ad - all species detected. Species other than those included in the input files are grouped together as ‘other’ in a gray stripe at the top of the Ad bar. QIIME/UCLUST bars represent genus-level assignments.

#### QIIME/UCLUST

QIIME is a highly popular metagenomics analysis program that was developed to analyze reads generated by 16S rRNA gene amplicon sequencing rather than full metagenome shotgun sequencing data (12). To accommodate this, we used bowtie2 to select the reads from our *in silico* communities that matched 16S rRNA genes in the GreenGenes v13.8 database and created new input fastq files containing only those reads, a protocol that has been previously used to enable QIIME analysis of ancient metagenomic sequences (8). The taxonomic proportions of the 16S rRNA gene input files were initially skewed by the bowtie2 identification such that some taxa were over-represented while others were under-represented relative to the full genome proportions (Fig. 4, S9-S10, bars If *vs.* 16f, Supplemental Table S3). As the 16S rRNA gene does not provide species-level resolution for many species, we assessed the accuracy of assignments at the genus level. QIIME/UCLUST failed to identify 2, 17, and 19 input taxa in the 40-, 100-, and 200-genome simulated datasets (22 total input taxa comprising 16 genera) (Fig. 3, S7-S8, Supplemental Table S4), despite the presence of reads derived from these 22 genomes in the bowtie2 16S rRNA gene-identified reads files. Of the missing taxa, 11 are not included in the GreenGenes v.13.8 database at the species or genus level.

QIIME/UCLUST identified the highest proportion of false-positive taxonomic assignments (Fig. 4, S9-S10) (“other” in barchart figure), and the proportion of false-positive taxa was higher in ancient than modern simulated datasets, suggesting that damage patterns decrease the accuracy of taxonomic identification by this program. Because of the large number of false-positive taxa identified, as well as the several taxa remaining unidentified, many of the input taxa were under-represented in the OTU tables produced by QIIME (Fig. 3, 4, S7-S8). Circular trees generated in metacodeR representing the taxonomy of the OTUs identified in the 40-genome ancient dataset full and filtered table taxonomic assignments (Fig. 5, S19) show that QIIME/UCLUST tends to overestimate each phylum in proportion to the original input, except for poorly characterized taxa such as Candidate divisions TM7 (Candidatus *Saccharibacterium)* and SR1, *Spirochaetes*, and archaea, and there is a slight bias toward over-assignment of *Proteobacteria*.

**Figure 5.**
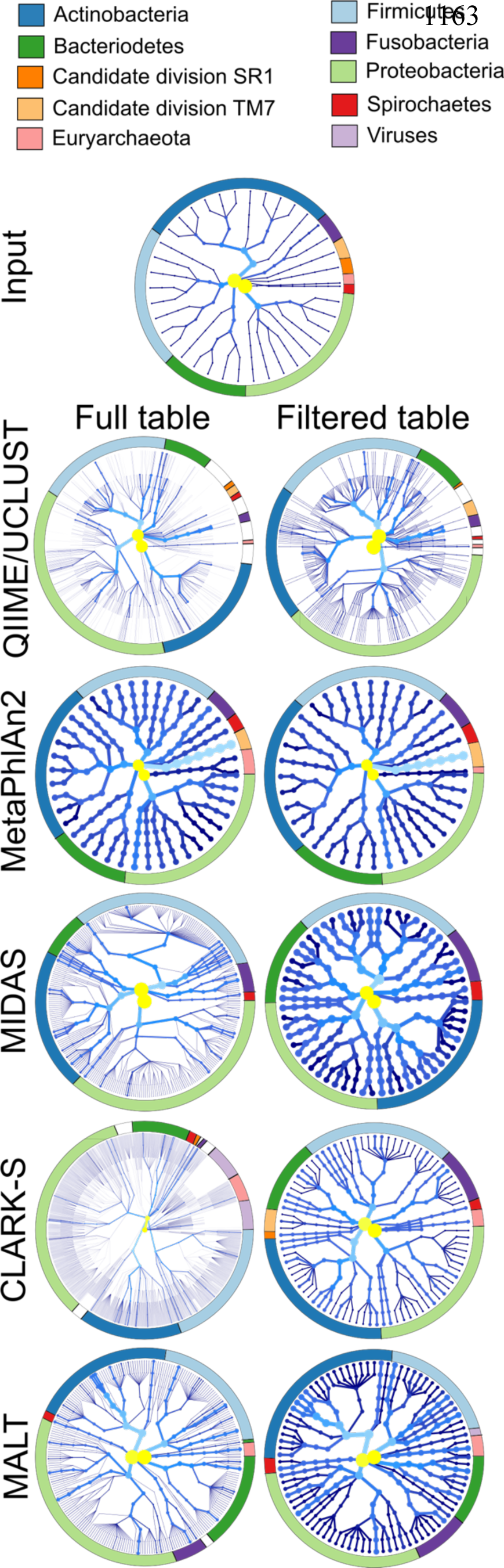
Biases in species detection across the phylogenetic tree are database-dependent. Species detected by each program represented in a radial phylogenetic tree with the nodes representing different taxonomic levels, where innermost node is root and the outermost nodes are strains. More highly represented taxa are lighter in color (yellow to light blue) and have thicker branches/nodes, while less abundant taxa are darker blues with thinner branches/nodes. The ring encircling each tree designates the major phyla (those in the input files, plus viruses when distinguishable) by color. For programs that did not report strains (QIIME/UCLUST, MetaPhlAn2, CLARK-S, MALT) the species was repeated as a strain to maintain consistency with MIDAS.

We identified several genus-level false-positive taxa with particularly high assignments in the 40-, 100-, and 200-genome datasets individually, as well as 7 that were shared by all 3 datasets (Supplemental Table S4). Three of the 7 genera, *Bacteriodes, Coprococcus*, and *Enterococcus*, had high numbers of assignments only in the ancient simulated datasets, while *Achromobacter, Actinobacillus, Enterobacter*, and *Erwinia* were highly represented in both ancient and modern simulated datasets. The genomes from which the reads assigned to each of these 7 false-positive taxa originated were identified (Supplemental Table S4), and we tested if these assignment biases hold true in real datasets. All reads assigned to the 7 false-positive genera in set of historic calculus samples from the Radcliffe Infirmary burial ground (ca. 1770-1855; Oxford, England) (3) were searched against the NCBI nt database using BLASTn to identify the likely species of origin for the reads. Many of the biases in read assignment observed in the *in silico* datasets were also observed in the real calculus samples (Table S4), i.e., *in* silico-generated reads assigned to *Enterococcus* by QIIME were from taxa in the order *Lactobacialles*, and historic calculus reads assigned to *Enterococcus* by QIIME also had best BLAST hits to the order *Lactobacialles.*

#### DADA2

DADA2 (26) and deblur (28) are new methods for taxonomic assignment of 16S rRNA gene reads that have been implemented in QIIME v2.0. Rather than using a percent similarity cut-off for assignment of a read to an operational taxonomic unit, these programs use exact sequence matches, and rely on Illumina sequencing error models to determine if single nucleotide polymorphisms in a read are true sequence variation or the product of sequencing error. The implementation of these programs requires multiple copies of each sequence, which while common in 16S rRNA gene amplicon datasets, are not likely to occur in a set of ancient reads that are selected out of a shotgun sequenced metagenomic dataset, such as we performed, due to insufficient coverage. As a result, we were unable to run DADA2 through to taxonomic assignment of our 16S rRNA gene-identified reads because each read was represented only once in each of our datasets. Because 16S rRNA gene amplification from ancient DNA samples has been shown to produce strong taxonomic biases (8), and because DADA2 is unable to classify the low coverage reads typical of shotgun metagenomic datasets, we recommend against using QIIME v2.0 for taxonomic characterization of ancient microbial samples, and any low-coverage non-amplicon data.

#### MetaPhlAn2

MetaPhlAn2 is a fast program that assigns taxonomy based on single-copy marker genes that are unique to each species in the MetaPhlAn2 database (14, 15). It was shown to be highly accurate for assigning taxonomy in modern metagenomic samples, and it is implemented in metaBIT (29), a user-friendly wrapper program that is targeted to ancient metagenomics researchers. MetaPhlAn2 identified the smallest number of false-positive taxa of the programs tested (Supplemental Table S5), but had exceptionally skewed proportions of 2 identified taxa, which may explain why weighted UniFrac distance community structures were so different from truth, while unweighted UniFrac distance community structures were highly similar to truth, where truth represents the percent of DNA from a genome rather than the cell count. Circular taxonomic assignment trees of the ancient dataset demonstrate that MetaPhlan2 does not report high numbers of false-positive taxa in any phylum (Fig. 5, S19), and although there are 3 more *Proteobacteria* reported than in the input files, they were identified at low-abundance and removed during filtering. The only phylum not represented in the MetaPhlAn2 output dataset is Candidate division SR1, which is not in its database.

Candidatus *Saccharibacterium* TM7b was represented at 1500-2000% higher relative abundance in the output files than the abundance of DNA in the input files in 40-, 100-, and 200-genome datasets, both ancient and modern (Fig. 3, 4, S7-S8, S11-S12). This may be the result of the MetaPhlAn2 normalization method, which calculates the proportion of cells from each species based on single-copy marker genes, rather than reporting the relative abundance of all DNA from each species detected (14). The TM7 genome is much smaller than the genomes of the other species we included in our dataset, 0.1 Mb *vs.* 2.5-3.5 Mb, and because our datasets have the same number of reads from each species, there must be more copies of the small genome-cells in the datasets to achieve the same proportion of DNA. We calculated that our datasets have approximately 7.8 copies of the TM7 genome but on average 0.36 copies of all other species genomes, which is a difference of ~2000% (Supplemental Table S1). *Desulfobulbus* sp. oral taxon 041 was identified at 200-300% higher relative abundance in all output files, and *Prevotella* sp. oral taxon 299, present only in the 200-genome datasets, was identified at 200-300% higher relative abundance in the output files (Figs. 3, 4, S11-S12). Both of these organisms have small genomes, ~0.7 Mb, and like TM7b they have more cell copies per dataset than the average (1.2 *vs.* 0.36), which is a difference of ~340%.

Twenty-three input taxa were not specifically identified in any of the simulated datasets, and all were missing from the MetaPhlAn2 database or not present at the appropriate taxonomic level (Supplemental Table S5). Nine of the missing 22 taxa are subspecies, including four of *Fusobacterium nucleatum*, one of *Mycobacterium avium*, and 4 of *Salmonella enterica* subsp. *Enterica* (Supplemental Table S5), and these were identified to the species level (or in the case of *Salmonella enterica* to subspecies) but not lower. Several species are indistinguishable by the marker genes used by MetaPhlAn2, and are grouped together, including *Streptococcus mitis/oralis, Bordetella bronchiseptica/parapertussis*, and *Mycobacterium tuberculosis* complex *(tuberculosis/bovis/canetti/africanum)*. If a user wishes to specifically identify any of these species, other programs will need to be used. The 40-genome dataset had 7 false-positive taxa, the 100-genome dataset had 12, and the 200-genome dataset had 14, yet all were low abundance, suggesting that MetaPhlAn2 may be slightly less accurate making assignments in samples with higher diversity, and may minimally inflate that diversity. Only one false-positive taxon, *Streptococcus tigurinus*, was common to the 40-, 100-, and 200-genome datasets, and this may be because of inconsistencies in naming this *Streptococcus* species, where some NCBI entries use *tigurinus* as an independent species and others use it as a subspecies of *S. oralis.* The reads assigned to S. *tigurinus* may be from *S. oralis subp. tigurinis*, which was one of the input genomes we used.

#### MIDAS

MIDAS is another fast program that uses a panel of 15 single-copy marker genes present in all of the species included in its database to perform taxonomic classification (16). It also has the ability to determine differences in gene presence/absence and detect single nucleotide polymorphisms (SNPs), although these were not tested in this study. MIDAS has a substantial database (~31000 genomes) in which related species are grouped together under a single species identifier number (5952 total identifiers), which we found introduces biases in the species reported in the output tables. A majority of the species detected in each dataset were found only in ancient samples (82%, 72%, 64% in the 40-, 100-, and 200-genome datasets, respectively), yet these had a relative abundance of <0.1%, suggesting that aDNA damage patterns do lead to false assignments in MIDAS, but only of a small number of reads. *Streptococcus* species were the most common low-abundance false-positive taxa, and likely indicate a database bias. Biases in reporting *Firmicutes* and *Proteobacteria*, and to a lesser extent *Actinobacteria*, in ancient datasets can be seen in the circular taxonomic assignment trees (Fig. 5, S19). Filtering low abundance hits removes many of the false-positive taxa in these phyla, yet several lower-abundance (darker nodes/branches) false-positive taxa remain in each. MIDAS did not report any archaea, despite having the input species in the database, nor does it detect Candidate divisions TM7 and SR1, which are not in the database.

In total, MIDAS failed to identify 28 input taxa, the highest number missed of all the programs we tested, and only 3 of these missed taxa were not in the database (Supplemental Table 6, Figs. 3, S7-S8). Despite missing so many species, MIDAS maintained relative proportions of the input taxa in even species distribution, and the proportion of false-positive taxa detected was slightly lower in ancient simulated datasets than modern (Fig. 4, S13-S14). To understand why we saw certain abundant false-positive taxa, we investigated the origin of the reads assigned to five false-positive taxa that were highly abundant in the 40-, 100-, and 200-genome datasets. We determined which, if any, additional species shared the same MIDAS-specific species identifier number, and the origin of the reads being assigned to these false-positive taxa (Supplemental Table S6). Reads from several taxa not reported by MIDAS were assigned to false-positive taxa, explaining both why certain taxa were missed and the high abundance of these false-positive taxa.

This phenomenon highlights how grouping related species under a single species identifier and presenting only one of those species in the output table can result in curious species profiles from MIDAS. For example, *Phocaeicola abscessus*, which had high abundance in 100 and 200-genome datasets but was not part of the input files, shares an identifier number with *Bacteriodetes* oral taxon 272, which was in the input files but was absent from the final species tables MIDAS produced. By checking the alignment files that MIDAS generates, we determined that the reads from *Bacteriodetes* oral taxon 272 were assigned to the species identifier shared by these two organisms. The same was true for other false-positive taxa/missing taxa pairs including *Actinobaculum* sp. and *Actinobaculum* sp. oral taxon 183, *Bordetella bronchiseptica* and *B. pertussis*/*B. parapertussis*, S*ynergistetes* bacterium and *Fretibacterium fastidiosum, Fusobacterium nucleatum* CC53 and *Fusobacterium nucleatum* subsp. *vincentii*, and Candidatus *Prevotella* and *Prevotella* oral taxon 317 (Supplemental Table S6). Most of these biases are against oral taxa, which is not surprising for a program developed using non-oral microbiome sources. MIDAS also has difficulty making assignments to the genera *Neisseria, Fusobacterium*, and *Salmonella* (Supplemental Table S6), and slightly overestimates *M. tuberculosis* and *Y. pestis* (Figs. S7-S8), suggesting a slight bias for potentially human pathogenic organisms.

#### CLARK-S

CLARK-S, a version of the CLARK sequence classification system (19, 20), uses spaced k-mers to match reads to whole genomes in a database, and was developed specifically to classify reads in metagenomics samples. It performs similarly to Kraken (18), makes assignments only at the taxonomic level designated by the user (default species), and cannot report strains or sub-species. As the database size for CLARK-S increases, the amount of memory required to generate and load the hash table increases substantially, and our database of 16855 genomes required 1TB of memory (necessitating use of a high-performance computing cluster), yet the program classified each sample in a few hours. CLARK-S was the only program that detected all of the species in the input files that were in the database (Supplemental Table S7) (all of the genomes used to create the input files were deliberately included in the CLARK-S custom database); however, it also reported the highest number of false-positive taxa (~6000 in each 40-, 100-, and 200-genome dataset).

A majority of the species detected were present only in ancient simulated datasets (80%, 75%, and 69% of species in 40-, 100-, and 200-genome datasets, respectively), and the overwhelming majority were present at <0.1% abundance. As filtering all species with relative abundance <0.1% removed most of the low-abundance false-positive taxa but only 1-2% of the total assigned reads, we recommend filtering all tables generated by CLARK-S. There was no clear distinction between high and low abundance false-positive taxa, unlike in several other programs we tested. Instead there was a steady decrease in the abundance of false-positive taxa, with a very long tail of very low abundance species.

Circular taxonomic assignment trees of the CLARK-S unfiltered tables show slight biases for *Actinobacteria*, but mostly overestimate each phylum in proportion to the original input (Fig. 5, S19). A substantial number of viruses were reported, but were all reported at <0.1% abundance and removed by filtering. Most of the input species were detected by CLARK-S at proportions close to those of the input files for 40-, 100-, and 200-genome datasets, both ancient and modern (Fig. 4, S15-S16), but it was poorly able to detect the genera *Bordetella, Burkholderia, Mycobacterium*, and *Yersinia* (Fig. 3, S7-S8). Generally, species overestimation was lower in the modern than ancient samples, but underestimation was not consistently different between ancient and modern sample sets (Fig. 3, S7-S8).

#### MALT

Like CLARK-S, MALT (21) uses spaced hashes to classify reads to the genomes in a database, and it is the only program we tested that can align reads to a protein database, done through BLASTx, which also allows functional characterization of the microbial community. We ran MALT in BLASTx-mode (22) to assess how translating the ancient simulated metagenomic reads affected taxonomic profiles, using a database consisting of NCBI RefSeq non-redundant bacteria, viral, archaeal, and plasmid protein sequences (57435 species/strains). The amount of memory required to load the hash table into memory was >1TB (again necessitating use of a high-performance computing cluster), and the program classified samples more slowly than CLARK-S, requiring several hours longer per sample than CLARK-S. The output files were uploaded to MEGAN6 (30, 31) and read count and relative abundance tables of only species-level assignments were exported, although MALT does place reads higher up on the taxonomic tree if they cannot be assigned to a species with high confidence. Fourteen input taxa were missing from the output files, 9 of which were not in the database (Supplemental Table S8). However, reads from each of these taxa were assigned to higher taxonomic levels, and in low numbers to closely-related species that were not in the input files.

MALT overestimated the number of species in all datasets, but the difference in the total number of assignments between ancient and modern datasets was much smaller than CLARK-S (Fig. 2C, S5C-S6C). Circular taxonomic assignment trees show a bias for *Proteobacteria* that remains after filtering (Fig. 5, S19). In the ancient simulated datasets there were 54, 86, and 75 species detected in the 40-, 100-, and 200-genome datasets, respectively, that were not reported in the modern datasets. The over/underestimation of the relative abundance of each input species was consistent between modern and ancient samples (Fig. 3, 4, S7-S8, S17-S18). The 40-, 100-, and 200-genome datasets each had 5 had false-positive taxa present at >0.1% abundance, while two of these false-positive taxa were reported in the all 3 genome datasets. We observed that MALT assigned a low number of reads to a particularly high number of *Neisseria* and *Prevotella* species that were not in the input files. In the 40-, 100-, and 200-genome datasets, MALT identified 32, 19, and 17 false-positive *Neisseria* species, respectively, and 22, 37, and 34 false-positive *Prevotella* species, respectively, although all of these species were present at <0.1% abundance. This may be because the number of species in the database from these genera is higher than for other species in the input files (such as *Actinobacteria* and *Fusobacteria).*

One unusual false-positive taxon that was consistent with MIDAS was *Phocaeicola abscessus* in the 100 and 200-genome datasets, both ancient and modern, at a relative abundance of 0.4-0.9%. The reads assigned to *P. abscessus* were all from the *Bacteroides* sp. oral taxon 272 genome, and *Bacteroides* sp. oral taxon 272 was identified at approximately 10% lower relative abundance than *P. abscessus* in all samples. Candidatus *Saccharibacterium* oral taxon TM7x had high numbers of reads assigned to it despite not being in the input file, but it was the only Candidatus *Saccharibacterium* TM7 species in the database and the reads assigned to it were from the TM7 genomes included in the input files. MALT classified a very small number of reads per sample to viruses (<50), but the assignments were not to species level, and were not included in the output files we analyzed.

## Discussion

Reconstructing microbial community composition and structure from short sequencing reads is challenging (32), especially from highly damaged ancient DNA data-sets. Here we show that biases inherent to specific taxonomic assignment programs are more pronounced than biases arising from ancient DNA damage patterns. Each program we tested has intrinsic, and at times non-intuitive, assignment biases, and an appreciation of these biases is needed to aid interpretation and limit inappropriate conclusions.

Our study does not show that one program clearly outperforms others, but rather each has unique advantages and disadvantages. For example, for accurate interpretation of community structure, MIDAS is an appropriate choice if species relative abundance is critical (ie, by weighted UniFrac distance), while MetaPhlAn2 is more appropriate if relative abundance is not critical (ie, by standard UniFrac distance). However, taxonomic accuracy in MIDAS is hampered by the way that the species are reported. For example, while MIDAS reduces potential assignments from tens of thousands of genomes in its full database to a more manageable 5952 ID clusters that are actually used at the taxonomic assignment step, and it reports as the identified species for each query sequence only one representative species per ID cluster, resulting in inappropriate species profiles despite reads being assigned to an appropriate genome. It may be possible to correct this effect by altering the program to preferentially select a different representative species appropriate for the sample type under analysis, but this would require alteration of the source code or substantial reanalysis of the output files.

One major difference between the different programs test here lies in the way these compute relative abundance. By using a set of single-copy marker genes, both MetaPhlAn2 and MIDAS attempt to report the proportion of cells of each species detected in a sample. This is in contrast to k-mer-based methods such as CLARK-S and MALT, which report the proportion of total DNA assigned to each species. This difference may explain why the community structures (beta-diversity) reported by MetaPhlAn2 and MIDAS were closest to the simulated values. Genome size can vary substantially between bacterial species, and those with larger genomes may appear more abundant in a sample because a higher proportion of DNA is from those species, even though the number of cells may not be higher. Species relative abundance reported by k-mer-based identification methods can be normalized by predicted genome size in order to approximate cell copy number even when the exact strain is not known, as genome size is largely consistent within species. The distinction between the relative abundance reported by cell copy-normalizing (MetaPhlAn2 and MIDAS) and non-normalizing (CLARK-S, MALT, QIIME) metagenomic profilers should be kept in mind when considering metagenomic community profiles.

For maximizing the number of assigned reads or determining the relative abundance of all DNA fragments CLARK-S is best (if, for example, one wants to attempt genome assembly from all reads assigned to a species). Detecting genuine low-abundance species, however, especially viruses and bacteriophages, cannot be achieved with CLARK-S due to a high rate of false-positive identification with abundance lower than 0.1%. MALT is unique in that it can provide functional classification of reads as well as taxonomic classification, but it has difficulty making assignments when the database used has a high number of closely-related species (discussed below). In addition, similarly to CLARK-S, MALT has a high rate of false-positive assignment at low abundance. QIIME/UCLUST provides the least accurate method, which included many false positives even when low-abundance taxa were filtered out. In addition, our results indicate that it is the only program whose performance was distinctly different between ancient and modern samples, and the differences could not be resolved by removing low-abundance taxa.

Most of the program-specific biases we observed were due to the database each program used. Familiarity with the taxa present in modern samples is important to ensure appropriate species representation in the database being used, and to customize the databases when possible. This will be much more straightforward for relatively well-characterized human body sites such as the mouth (4), and to a lesser extent the gut (33), but will be more nuanced for poorly characterized communities such as those from non-model organisms (34–36). For example, the default RefSeq bacteria database downloaded by CLARK-S does not include any species of *Actinomyces*, and has very few species of *Prevotella*, both of which are prevalent and highly-abundant oral genera, and the latter of which is major taxon in the gut microbiota of traditional societies (37). Restricting the database to RefSeq genomes alone, such as we did for MALT, limits the genomes to those that have been quality-checked and curated, and most sequenced genomes have not met these criteria, nor have metagenome-assembled genomes. Finally, the GreenGenes taxonomy has not been updated since 2013 and contains now-obsolete taxonomic classification for some organisms, which can confuse results, and more recently updated taxonomic classification systems (38) should be used.

Although ancient dental calculus is highly resistant to taphonomic processes and infiltration of environmental contaminants, it is not immune from these processes, and palaeofeces and other non-calcified archaeological specimens (39) are particularly susceptible to environmental contamination and degradation. Environmental microbes, particularly from soil burial matrix and skin of individuals handling the samples, may remain associated with archaeological samples after cleaning and sterilization and contribute to the metagenomic profile generated by sequencing. Distinguishing environmental signatures from endogenous signatures will be critical for ensuring accurate reconstruction of host-associated microbial profiles. Although outside the scope of this discussion, most microbial databases are heavily dominated by human-associated bacteria, and this may bias the assignment of soil and environmental species. Approaches for limiting false identification of environmental microbial species as host-associated species are discussed in Warinner, *et al.* (11).

The simulated ancient metagenomic datasets we generated were modeled after data generated from archaeological dental calculus (3), and we selected 5M reads for the *in silico* samples because this was the lowest read count in these samples. However, McIntyre, *et al.* 2017 (25) have shown that as read depth increases the performance of metagenomic classifier tools changes, and this should be kept in mind for studies with higher sequencing depth. We chose not to normalize the output from each program to a consistent taxonomic level, such as genus, because we wanted to work with data that was as close to the default output as possible. This allowed us to see the resolution limit of the programs with respect to known species, subspecies, and strains, as well as the strengths and weaknesses of that resolution. While higher taxonomic classification may demonstrate broad community level changes, the immense genetic variation in strains of a single bacterial species, for example *Streptococcus mutans* (40), prevents accurate prediction of changes in metabolic functional capacity from higher order taxonomy.

It is important to note, however, that while community resolution is lost when reads are assigned to higher levels of taxonomy, this technique may ultimately retain more information. Community structure may be better estimated at levels of taxonomy higher than species because reads that do not have species-level resolution can be classified at higher taxonomic levels with greater confidence. Using an LCA (lowest common ancestor) algorithm, MALT assigns reads to higher taxonomic levels if they cannot be distinguished between two nearly-genetically identical species. For example, some species within the genera *Yersinia* (*Y. pestis* and *Y. pseudotuberculosis*) and *Bordetella* (*B. pertussis, B. parapertussis, B. bronchiseptica*) are highly genetically similar, and reads that map equally well to multiple species in those genera are usually assigned at the genus level by the LCA algorithm in MALT. Similarly, QIIME/UCLUST will classify reads to deeper nodes in the tree by if they cannot be assigned to lower taxonomic levels. For example, the percent of reads in our dataset assigned to different taxonomic levels were: species – 17%, genus – 65%, family – 13%, order – 2.4%, and class – 0.7%. Users should be aware of this behavior in specific programs, and be aware of the node at which reads from those taxa tend to assign, as this can substantially affect analyses performed only at the species level.

Assigning taxonomy to reads below species level is desirable to understand the functional capacity of the microbial community, but the programs we tested performed this task poorly. The ability of MIDAS to discriminate strains or subspecies varies considerably by organism. For example, the 12 strains of *Porphyromonas gingivalis* in the database share the same species ID, while the 31 strains of *Streptococcus mitis* each have a unique species ID. This resulted in the MIDAS-produced species profiles containing one strain of *P. gingivalis* in the 100 and 200-genome datasets (despite there being two and four, respectively), and 29-30 strains of *S. mitis* across each 40-, 100-, and 200-genome dataset (albeit all very low abundance), despite there being only one species in all three datasets. To avoid biases of strain-level identification by this program, we combined all strain-level assignments of the same species into one species-level assignment for all analyses If identifying subspecies or strains present in a sample is desired, programs specifically designed to perform this function, such as StrainPhlAn (41), Sigma (42), or Platypus Conquistador (43), are recommended instead. Furthermore, special care should be taken to ensure results are not false positives or derived from modern environmental contamination by following guidelines suggested by Warinner *et al.* (11) and Key, *et al.* (44).

High proportions of the fecal-associated genera *Coprococcus, Enterococcus*, and *Enterobacter* were identified by QIIME/UCLUST in our *in silico* generated dataset, but they were not in the input files. Rather, a high number of reads of consistent taxonomy were assigned to these genera, which we confirmed occurs in real datasets, indicating that these assignments are more likely an artifact of the taxonomic classification process than an indication of poor hygiene. This demonstrates how interpreting taxonomic assignment results without understanding the biases and limitations of the program used could lead to erroneous conclusions about microbial community profiles, and ultimately human activity.

Identifying bacteriophage in ancient metagenomic samples is challenging and new methods are needed. MetaPhlAn2, CLARK-S, and MALT all detected phages in very low abundance, below levels of suggested filtering to remove spurious assignments. Active bacteriophage replication in the oral biofilm is associated with altered host health status (5, 45), and monitoring phage activity may offer insight into biofilm pathogenicity in oral (45, 46) and gut (47) sites. Therefore, reliably detecting bacteriophage in host-associated ancient metagenomic samples may allow us to study phage-mediated biofilm changes and evolution relating to human disease. While it is unlikely that we will be able to determine if phage-identified sequencing reads are from viral particles or host-integrated prophages, proteomic characterization of ancient microbiomes (1) may be able to detect viral proteins indicating free phage particles.

Recently, McIntyre, *et al.* (25) assessed performance of a wide selection of metagenomic taxonomic classification programs built upon a variety of techniques. They reported that the precision of taxonomic assignment can be improved by combining results of certain programs that use different assignment methods, including MetaPhlAn2 and CLARK-S. Combining the results of these taxonomic assignment programs for ancient metagenomics samples may then increase reliability and confidence in historic community structure and composition, and should be examined further with *in silico*-generated datasets. Confirming species presence/absence by detection with two independent taxonomic classifiers will assist with ensuring specific program biases are not reported as true results.

There are several factors that we did not test that may influence taxonomic profiling of ancient DNA. These include environmental contamination (11) (discussed above) sample location- and age-specific differences in damage patterns (48), and species-specific differential preservation of bacterial DNA (8). Additional *in silico* dataset testing, such as by using mapDamage profiles modeled after older archaeological samples, samples from different locations, or based on reads mapped to different or multiple species, may be warranted to determine if and how strongly these factors affect taxonomic profiling. Based on our results that age-related damage patterns minimally affect read taxonomic assignment, however, we do not expect these variables to substantially alter taxonomic profiles. Nevertheless, location- and age-related biases should be considered in studies that compare samples across geographic locations and/or time.

We have demonstrated that the damage patterns characteristic of ancient DNA do not substantially affect taxonomic profiling by the five programs we tested. Instead the biases we detected are inherent to the programs themselves and the database each program uses. This is promising for comparing ancient microbiome samples with modern samples when using the same taxonomic classifier, as biases will be shared by both. Our results highlight the importance of knowing the limitations of the metagenomic classifier being used, and investigating any unusual results, such as the presence of unexpected taxa and the absence of expected taxa, to ensure appropriate interpretation of taxonomic profiles.

## Materials and Methods

### Simulated ancient and modern metagenomics samples

Simulated ancient and modern metagenomics fastq files were generated with gargammel (27). Samples of 5 million reads, 99% bacterial and 1% human were generated with 40-genomes, 100-genomes, or 200-genomes, with even genome distribution (equivalent numbers of reads from each input genome), and both with and without simulated ancient DNA damage patterns, and sequencing errors were based on Illumina HiSeq2500 150bp paired-end chemistry and default Illumina adapters. Thirty-nine total metagenomes were simulated as follows: 40-genome even distribution ancient (10) and modern (3), 100-genome even distribution ancient (10) and modern (3), and 200-genome even distribution ancient (10) and modern (3). Genomes are listed in

Supplemental Table S1, and were selected to resemble dental plaque bacterial communities based on the species listed in the Human Oral Microbiome Database (homd.org), and relative abundance was roughly based on dental plaque-derived biofilm composition (Velsko & Shaddox, in review). Select non-oral bacterial species were added to assess biases in detecting specific “pathogenic” species. Although the genomes are represented with equal proportions of DNA in each dataset, the number of cells from each organism is unevenly distributed because of differences in genome size (Supplemental Table S1).

Age-related damage patterns were simulated based on mapDamage (9, 10) base composition file and misincorporation file generated on analysis of real historic dental calculus metagenomic samples sequenced on an Illumina HiSeq2500 with 150bp paired-end chemistry, with bacterial genome damage patterns based on reads mapped to the Tannerella forsythia 92A2 genome (assembly GCA_000238215.1) (Fig. S1) and human genome damage patterns based on reads mapped to the human genome (assembly GCA_000001405.26) (Fig. S1), while the fragment length distribution was based on all reads in sample CS21. Simulations for modern metagenomics samples did not include damage pattern input files. The command to simulate ancient metagenomic samples was: ./gargammel.pl --comp 0.99,0,0.01 -n 5000000 --misince dnacompCS32e.txt --misincb dnacompCS21b.txt -f fragmentlengthCS21.txt -mapdamagee misincorporationCS32.txt single -mapdamageb misincorporationCS21.txt single -rl 150 -ss HS25 -o output/anc40e1 input/. The command to simulate modern metagenomic samples was: ./gargammel.pl --comp 0.99,0,0.01 -n 5000000 -l 150 -rl 150 -ss HS25 -o output/mod40e1 input/. Damage profiles for human (CS21) and bacterial (CS32) reads came from different calculus samples because these had the highest number of reads to the human and *T. forsythia* genomes, respectively, which allows the most accurate assessment of damage profiles (11). Fragment length distribution for ancient simulated samples was based calculus sample CS21, while read length of 150bp was specified for modern samples. The genome of origin for each read is included in the read name by a gargammel-generated code (listed in Supplemental Table S1), and the exact number of reads derived from each genome was determined by counting in each of the 78 input fastq files.

### Read processing and 16S rRNA gene fragment filtering

Reads were processed following a custom pipeline optimized for ancient DNA metagenomics samples. AdapterRemoval (49) was used to detect and remove consensus adapter sequences, quality-trim reads at Q30 and collapse paired reads. Singleton files were discarded and reads with residual adapters were detected with bowtie2 (50) and filtered from the samples with filter_fasta.py in QIIME v1.9 (12). Four final files were generated: collapsed reads, pair 1 reads, pair2 reads, and truncated collapsed reads, and all 4 files were concatenated to generate a single input file for taxonomic classification. Reads mapping to the 16S rRNA gene were identified in the independent final files and collected in separate files for classification as follows. A bowtie2 database was generated from the GreenGenes v13.8 database (51), and the cleaned and collapsed, pair1, pair2, and collapsed truncated fastq files were searched against this database with bowtie2. All reads that mapped to 16S rRNA gene reads were filtered from the full fastq files to a separate file using seqtk (https://github.com/lh3/seqtk). All 4 files matching the 16S rRNA gene (collapsed, pair1, pair2, and collapsed truncated) were concatenated for taxonomic classification (Supplemental Table S3).

### Taxonomic classification

Reads in all simulated metagenomics samples were classified with 5 taxonomic identification programs (Supplemental Table S2): QIIME v1.9/UCLUST/GreenGenes v13.8 database (12, 51, 52), MetaPhlAn2 (14, 15), MIDAS (16), CLARK-S (20), and MALT (21) run in BLAST-X mode (22). All options that differed from default are listed in Table S2. Each program uses a different classification method. QIIME v1.9 was used to bin reads matching the 16S rRNA gene using UCLUST (52) with pick_closed_reference_otus.py and to assign taxonomic classification with the GreenGenes v 13.8 database at 97% identity (202421 sequences, 99322 OTUs). Samples were not rarefied to identical OTU counts prior to analysis, as this practice has been shown to be unnecessary (53). The output biom file was summarized at the species level, which included all assignment levels kingdom through species. MetaPhlAn2 and MIDAS used their respective default databases (16904 species/strains, 31007 genomes/5952 species groups, respectively), while CLARK-S was run against a custom database of 16855 genomes, and MALT was run in BLAST-X mode against a custom database of NCBI RefSeq non-redundant bacteria, viral, archaeal, and plasmid protein sequences (57435 species/strains). MetaPhlan2 and CLARK-S output were set to species level. The MALT output rma6 files were uploaded to MEGAN6 (31), and classification tables of species assignments only were exported. Output for each classification program is unique, with MetaPhlAn2 and MIDAS providing relative abundance on a scale of 0–100 and 0–1, respectfully, QIIME and CLARK-S providing a read count table, and MALT providing both relative abundance and read counts.

Outputs were normalized in 2 ways, generating 2 sets of tables: relative abundance of all assignments on a scale of 0-100 was calculated based on the number of reads assigned, if provided (QIIME, CLARK-S, MALT), and pseudo read counts were determined by multiplying the relative abundance by the total number of reads in the input files for MetaPhlan2 and MIDAS. The true input tables were also converted to biom format in count read and relative abundance formats. The NCBI taxonomy ID of each taxonomic assignment in each program was determined and used to create a single taxonomy file to assign taxonomy to biom files. All output tables, read counts and relative abundance, were converted to biom format in QIIME v1.9 and taxonomy based on NCBI taxonomy ID was added to each. To determine if removing very low abundance assignments improved the profiles, a second set of biom files was generated by removing all assignments present at less than 0.1% abundance (filtered tables). All biom files were summarized at the phylum, class, and genus levels in QIIME v1.9 using summarize_taxa.py, to allow assessment of classification biases at different taxonomic levels. Mapping data, including the simulated age of the sample (ancient or modern) and the taxonomic assignment program, were added to the biom files in QIIME v1.9.

QIIME v1 is no longer being supported with the release of QIIME v2.0, and QIIME v2.0 uses different taxonomic assignment programs from QIIME v1: (DADA2 (26) and deblur (28)). We also tried to include DADA2 in this assessment (Supplemental Table S2), using 16S rRNA gene-identified reads and the DADA2 R package as follows. AdapterRemoval was run on the simulated samples as before, but pair1 and pair2 reads were not collapsed. The reads matching 16S rRNA genes were identified using bowtie2 and the GreenGenes v13.8 database as before and filtered out of the pair1 and pair2 files. The pair1 and pair2 files of 16S rRNA gene-identified reads were used as input in DADA2. DADA2 was not able to merge sequences in any file because all were unique, and this prevented DADA2 from performing the sequence variant calling, and the program was unable to perform taxonomic assignment. Therefore, we were unable to proceed with DADA2, and have no results to present.

### Diversity metrics

Alpha-diversity was calculated in QIIME using the metrics Faith’s phylogenetic distance, Shannon index, observed species, and Chao1, using count read and pseudo-count read files, and graphs were generated using Prism v7. Beta-diversity was calculated on relative abundance biom files and plotted using the R package phyloseq (54) for the metrics UniFrac (55) (accounts for phylogenetic relatedness and presence/absence) and weighted UniFrac (56) (accounts for phylogenetic relatedness, presence/absence and abundance), Bray-Curtis (accounts for presence/absence and abundance) and binary Jaccard (accounts only for presence/absence). A newick-formatted phylogenetic tree was generated with phyloT (http://phylot.biobyte.de) including the NCBI taxonomy IDs of all assignments made by each program (9919 total IDs), using the Internal nodes-Expanded and Polytomy-Yes options.

### Program assignment biases

All output text files were manually inspected for taxonomic assignment biases. When a species in the input files was not detected by a program, the database of that program was searched for that species to understand why it was missed. The percent over/under representation of each genome compared to the input file was calculated (relative abundance in output/relative abundance in input * 100) and plotted as a heat map with the R library gplots (http://www.rdocumentation.org/packages/gplots). The percent of each species in the input files detected by each program as well as the percent of all other species detected but not in the input file was plotted in R using ggplot2 (ggplot2.org). The R package metacodeR (57) was used to visualize phylogenetic tree assignment biases in the ancient datasets by each program. Each node is a taxonomic assignment starting with the root (yellow circle), then kingdoms, phyla, etc radiating off, to sub-species level at the tips. For programs that did not produce sub-species or strain-level taxonomic assignments, the species assignment was repeated, so maintain visual consistency between all trees. The input data for these trees is the species/subspecies level for all programs except QIIME/UCLUST (which included all levels), so the internal nodes sum the leaves moving from subspecies back towards the root. The colors and weight of nodes and branches represent the relative abundance of each taxonomic assignment, where lighter colors with thicker branches are more abundant (yellows and light blues) and darker, thinner branches are less abundant. The relative abundance is the average of all 10 output files for each program. A ring circling each tree and color-coding each phylum was added in Inkscape.

We tested whether a QIIME/UCLUST false-positive taxa read assignment bias was present in real ancient metagenomics samples by processing metagenomic data generated from 19th century dental calculus samples (3) through the same 16S rRNA gene selection and QIIME/UCLUST OTU-picking, and then filtering out all reads assigned to the designated “false-positive” genera. These reads were searched against the NCBI nt database with BLAST using default parameters and the BLAST hits of the reads assigned to each “false-positive” genus were determined using MEGAN6 and compared to the origin genomes of the false-positive-taxa assigned reads from the simulated samples.

## Data sharing and availability

All supplemental figures are available for download on figshare (as a single pdf): https://doi.org/10.6084/m9.figshare.5811285.v1

All supplemental tables are available for download on figshare (seperate tabs in a single excel spreadsheet): https://doi.org/10.6084/m9.figshare.5817837.v1

All gargammel-generated “raw” sequencing read files (forward and reverse) will be available for download when we find an appropriate site to host them. They’re big. We’re working on it. Please until then if you would like the files contact us and we’re happy to share.

## Acknowledgements

This work was supported by the Oxford University Fell Fund 143/108 (to G.L. and C.W.), the U.S. National Science Foundation (BCS-1516633 and BCS-1643318 to C.W.) and the U.S. National Institutes of Health National Institute of General Medical Sciences (2R01GM089886). L.A.F.F. was supported by a Junior Research Fellowship (Wolfson College, University of Oxford) and a Wellcome trust grant (210119/Z/18/Z).

We thank Dr. Louise Loe for access to the Radcliffe Infirmary burial ground collection. We thank James A. Fellows Yates and Dr. Krithivasan Sankaranarayanan for critical comments on the manuscript, and the Oxford Advanced Research Computing for use of the HPC cluster.

## Supplemental Tables and Figures

**Table S1.** Details of input metagenomic samples generated *in silico* by gargammel.

**Table S2.** Details of the 6 taxonomic classification programs used.

**Table S3.** 16S rRNA gene-identified read input file read counts per sample.

**Table S4.** QIIME/UCLUST-specific taxonomic assignment biases.

**Table S5.** MetaPhlAn2-specific taxonomic assignment biases.

**Table S6.** MIDAS-specific taxonomic assignment biases.

**Table S7.** CLARK-S-specific taxonomic assignment biases.

**Table S8.** MALT-specific taxonomic assignment biases.

**Fig. S1.** MapDamage plots showing damage patterns applied to bacterial reads (top panels, CS32 Tannerella forsythia reads) and to human reads (bottom panels, CS21 human reads). MapDamage plots are from real ancient dental calculus samples from ref. (3).

**Fig. S2.** Age-related damage patterns minimally influence reported non-phylogenetic-based community structure in 40-genome samples. Principal coordinates analysis plots of abundance-weighted Bray-Curtis distance and un-weighted Jacccard distance beta-diversity for datasets made with 40-genomes at the species and genus levels for full output tables and tables filtered to remove species present at < 0.1% abundance.

**Fig. S3.** Age-related damage patterns minimally influence reported non-phylogenetic-based community structure in 100-genome samples. Principal coordinates analysis plots of abundance-weighted Bray-Curtis distance and un-weighted Jacccard distance beta-diversity for datasets made with 100-genomes at the species and genus levels for full output tables and tables filtered to remove species present at < 0.1% abundance.

**Fig. S4.** Age-related damage patterns minimally influence reported non-phylogenetic-based community structure in 200-genome samples. Principal coordinates analysis plots of abundance-weighted Bray-Curtis distance and un-weighted Jacccard distance beta-diversity for datasets made with 200-genomes at the species and genus levels for full output tables and tables filtered to remove species present at < 0.1% abundance.

**Fig. S5.** Age-related damage patterns slightly increase within-sample diversity. Alpha diversity of 100-genome datasets calculated by (A) Faith’s phylogenetic distance, (B) Shannon index, (C) Observed species, and (D) Chao1 for full output tables and tables filtered to remove species present at < 0.1% abundance. MPA2 - MetaPhlAn2, anc – ancient simulated dataset, mod – modern simulated dataset.

**Fig. S6.** Age-related damage patterns slightly increase within-sample diversity. Alpha diversity of 200-genome datasets calculated by (A) Faith’s phylogenetic distance, (B) Shannon index, (C) Observed species, and (D) Chao1 for full output tables and tables filtered to remove species present at < 0.1% abundance. MPA2 - MetaPhlAn2, anc – ancient simulated dataset, mod – modern simulated dataset.

**Fig. S7.** Species detection and over/under-representation differ by program not age-related damage. Heat-map showing for each program tested the species relative abundance under-represented (blues), over-represented (yellows, oranges, reds), not detected (black), and accurately represented (white) relative to the true input files for modern and ancient 100-genome datasets. Where programs were unable to distinguish species, strains, or sub-species a single bar across those genomes is colored to represent the over/under-representation of the lowest identifiable taxonomic level. MPA2 - MetaPhlAn2, CLK-S - CLARK-S; A – ancient simulated dataset, M – modern simulated dataset.

**Fig. S8.** Species detection and over/under-representation differ by program not age-related damage. Heat-map showing for each program tested the species relative abundance under-represented (blues), over-represented (yellows, oranges, reds), not detected (black), and accurately represented (white) relative to the true input files for modern and ancient 200-genome datasets. Where programs were unable to distinguish species, strains, or sub-species a single bar across those genomes is colored to represent the over/under-representation of the lowest identifiable taxonomic level. MPA2 - MetaPhlAn2, CLK-S - CLARK-S; A – ancient simulated dataset, M – modern simulated dataset.

**Fig. S9.** QIME/UCLUST-specific differences in genus detection and relative abundance are consistent between ancient and modern 100-genome simulated datasets. Relative abundances of each bar represent: If - true input fasta file, 16f - 16S rRNA gene-identified read input fasta file (used for QIIME/UCLUST profiling), Id - input genera detected, and Ad - all genera detected. Genera other than those included in the input files are grouped together as ‘other’ in a stripe at the top of the Ad bar.

**Fig. S10.** QIME/UCLUST-specific differences in genus detection and relative abundance are consistent between ancient and modern 200-genome simulated datasets. Relative abundances of each bar represent: If - true input fasta file, 16f - 16S rRNA gene-identified read input fasta file (used for QIIME/UCLUST profiling), Id - input genera detected, and Ad - all genera detected. Genera other than those included in the input files are grouped together as ‘other’ in a stripe at the top of the Ad bar.

**Fig. S11.** MetaPhlAn2-specific differences in species detection and relative abundance are consistent between ancient and modern 100-genome simulated datasets. Relative abundances of each bar represent: If - true input fasta file, Id - input species detected, and Ad - all species detected. Species other than those included in the input files are grouped together as ‘other’ in a stripe at the top of the Ad bar.

**Fig. S12.** MetaPhlAn2-specific differences in species detection and relative abundance are consistent between ancient and modern 200-genome simulated datasets. Relative abundances of each bar represent: If - true input fasta file, Id - input species detected, and Ad - all species detected. Species other than those included in the input files are grouped together as ‘other’ in a stripe at the top of the Ad bar.

**Fig. S13.** MIDAS-specific differences in species detection and relative abundance are consistent between ancient and modern 100-genome simulated datasets. Relative abundances of each bar represent: If - true input fasta file, Id - input species detected, and Ad - all species detected. Species other than those included in the input files are grouped together as ‘other’ in a stripe at the top of the Ad bar.

**Fig. S14.** MIDAS-specific differences in species detection and relative abundance are consistent between ancient and modern 200-genome simulated datasets. Relative abundances of each bar represent: If - true input fasta file, Id - input species detected, and Ad - all species detected. Species other than those included in the input files are grouped together as ‘other’ in a stripe at the top of the Ad bar.

**Fig. S15.** CLARK-S-specific differences in species detection and relative abundance are consistent between ancient and modern 100-genome simulated datasets. Relative abundances of each bar represent: If - true input fasta file, Id - input species detected, and Ad - all species detected. Species other than those included in the input files are grouped together as ‘other’ in a stripe at the top of the Ad bar.

**Fig. S16.** CLARK-S-specific differences in species detection and relative abundance are consistent between ancient and modern 200-genome simulated datasets. Relative abundances of each bar represent: If - true input fasta file, Id - input species detected, and Ad - all species detected. Species other than those included in the input files are grouped together as ‘other’ in a stripe at the top of the Ad bar.

**Fig. S17.** MALT-specific differences in species detection and relative abundance are consistent between ancient and modern 100-genome simulated datasets. Relative abundances of each bar represent: If - true input fasta file, Id - input species detected, and Ad - all species detected. Species other than those included in the input files are grouped together as ‘other’ in a stripe at the top of the Ad bar.

**Fig. S18.** MALT-specific differences in species detection and relative abundance are consistent between ancient and modern 100-genome simulated datasets. Relative abundances of each bar represent: If - true input fasta file, Id - input species detected, and Ad - all species detected. Species other than those included in the input files are grouped together as ‘other’ in a stripe at the top of the Ad bar.

**Fig. S19.** Biases in species detection across the phylogenetic tree are database-dependent. Figure is identical to Fig. 5, but with a black background to better visualize color gradient and branch/node sizes of the trees. Species detected by each program represented in a radial phylogenetic tree with the innermost node as root and the outermost nodes as strains. More highly represented taxa are lighter in color (yellow to light blue) and have thicker branches/nodes, while less abundant taxa are darker blues with thinner branches/nodes. The ring encircling each tree designates the major phyla (those in the input files, plus viruses when distinguishable) by color. For programs that did not report strains (QIIME/UCLUST, MetaPhlAn2, CLARK-S, MALT) the species was repeated as a strain to maintain consistency with MIDAS.

## References

1. Warinner C, Rodrigues JFM, Vyas R, Trachsel C, Shved N, Grossmann J, Radini A, Hancock Y, Tito RY, Fiddyment S, Speller C, Hendy J, Charlton S, Luder HU, Salazar-García DC, Eppler E, Seiler R, Hansen LH, Castruita JAS, Barkow-Oesterreicher S, Teoh KY, Kelstrup CD, Olsen JV, Nanni P, Kawai T, Willerslev E, Mering von C, Lewis CM, Collins MJ, Gilbert MTP, Rühli F, Cappellini E. 2014. Pathogens and host immunity in the ancient human oral cavity. Nat Genet 46:336–344.

2. Warinner C, Hendy J, Speller C, Cappellini E, Fischer R, Trachsel C, Arneborg J, Lynnerup N, Craig OE, Swallow DM, Fotakis A, Christensen RJ, Olsen JV, Liebert A, Montalva N, Fiddyment S, Charlton S, Mackie M, Canci A, Bouwman A, Rühli F, Gilbert MTP, Collins MJ. 2014. Direct evidence of milk consumption from ancient human dental calculus. Sci Rep 4:7104.

3. Velsko IM, Overmyer KA, Speller C, Klaus L, Collins MJ, Loe L, Frantz LAF, Sankaranarayanan K, Lewis CM, Martinez JBR, Chaves E, Coon JJ, Larson G, Warinner C. 2017. The dental calculus metabolome in modern and historic samples. Metabolomics 13:134.

4. Dewhirst FE, Chen T, Izard J, Paster BJ, Tanner ACR, Yu WH, Lakshmanan A, Wade WG. 2010. The Human Oral Microbiome. J Bacteriol 192:5002–5017.

5. Ly M, Abeles SR, Boehm TK, Robles-Sikisaka R, Naidu M, Santiago-Rodriguez T, Pride DT. 2014. Altered oral viral ecology in association with periodontal disease. MBio 5:e01133–14.

6. Ghannoum MA, Jurevic RJ, Mukherjee PK, Cui F, Sikaroodi M, Naqvi A, Gillevet PM. 2010. Characterization of the Oral Fungal Microbiome (Mycobiome) in Healthy Individuals. PLoS Pathog 6:e1000713.

7. Bonner M, Amard V, Bar-Pinatel C, Charpentier F, Chatard J-M, Desmuyck Y, Ihler S, Rochet J-P, Roux de La Tribouille V, Saladin L, Verdy M, Gironès N, Fresno M, Santi-Rocca J. 2014. Detection of the amoeba Entamoeba gingivalis in periodontal pockets. Parasite 21:30.

8. Ziesemer KA, Mann AE, Sankaranarayanan K, Schroeder H, Ozga AT, Brandt BW, Zaura E, Waters-Rist A, Hoogland M, Salazar-García DC, Aldenderfer M, Speller C, Hendy J, Weston DA, MacDonald SJ, Thomas GH, Collins MJ, Lewis CM, Hofman C, Warinner C. 2015. Intrinsic challenges in ancient microbiome reconstruction using 16S rRNA gene amplification. Sci Rep 5:16498–19.

9. Ginolhac A, Rasmussen M, Gilbert MTP, Willerslev E, Orlando L. 2011. mapDamage: testing for damage patterns in ancient DNA sequences. Bioinformatics 27:2153–2155.

10. Jónsson H, Ginolhac A, Schubert M, Johnson PLF, Orlando L. 2013. mapDamage2.0: fast approximate Bayesian estimates of ancient DNA damage parameters. Bioinformatics 29:1682–1684.

11. Warinner C, Herbig A, Mann A, Fellows Yates JA, Weiss CL, Burbano HA, Orlando L, Krause J. 2017. A Robust Framework for Microbial Archaeology. Annu Rev Genomics Hum Genet 18:321–356.

12. Caporaso JG, Kuczynski J, Stombaugh J, Bittinger K, Bushman FD, Costello EK, Fierer N, Peña AG, Goodrich JK, Gordon JI, Huttley GA, Kelley ST, Knights D, Koenig JE, Ley RE, Lozupone CA, McDonald D, Muegge BD, Pirrung M, Reeder J, Sevinsky JR, Turnbaugh PJ, Walters WA, Widmann J, Yatsunenko T, Zaneveld J, Knight R. 2010. QIIME allows analysis of high-throughput community sequencing data. Nat Methods 7:335–336.

13. Schloss PD, Westcott SL, Ryabin T, Hall JR, Hartmann M, Hollister EB, Lesniewski RA, Oakley BB, Parks DH, Robinson CJ, Sahl JW, Stres B, Thallinger GG, Van Hor. DJ, Weber CF. 2009. Introducing mothur: open-source, platform-independent, community-supported software for describing and comparing microbial communities. Applied and Environmental Microbiology 75:7537–7541.

14. Segata N, Waldron L, Ballarini A, Narasimhan V, Jousson O, Huttenhower C. 2012. Metagenomic microbial community profiling using unique clade-specific marker genes. Nat Methods 9:811–814.

15. Truong DT, Franzosa EA, Tickle TL, Scholz M, Weingart G, Pasolli E, Tett A, Huttenhower C, Segata N. 2015. MetaPhlAn2 for enhanced metagenomic taxonomic profiling. Nat Methods 12:902–903.

16. Nayfach S, Rodriguez-Mueller B, Garud N, Pollard KS. 2016. An integrated metagenomics pipeline for strain profiling reveals novel patterns of bacterial transmission and biogeography. Genome Research 26:1612–1625.

17. Darling AE, Jospin G, Lowe E, Matsen FA, Bik HM, Eisen JA. 2014. PhyloSift: phylogenetic analysis of genomes and metagenomes. PeerJ 2:e243.

18. Wood DE, Salzberg SL. 2014. Kraken: ultrafast metagenomic sequence classification using exact alignments. Genome Biology 15:R46.

19. Ounit R, Wanamaker S, Close TJ, Lonardi S. 2015. CLARK: fast and accurate classification of metagenomic and genomic sequences using discriminative k-mers. BMC Genomics 2014 15:1 16:236.

20. Ounit R, Lonardi S. 2016. Higher classification sensitivity of short metagenomic reads with CLARK-S. Bioinformatics 32:3823–3825.

21. Vågene ÅJ, Herbig A, Campana MG, Robles García NM, Warinner C, Sabin S, Spyrou MA, Andrades Valtueña A, Huson D, Tuross N, Bos KI, Krause J. 2018. Salmonella enterica genomes from victims of a major sixteenth-century epidemic in Mexico. Nat Ecol Evol 1.

22. Weyrich LS, Duchene S, Soubrier J, Arriola L, Llamas B, Breen J, Morris AG, Alt KW, Caramelli D, Dresely V, Farrell M, Farrer AG, Francken M, Gully N, Haak W, Hardy K, Harvati K, Held P, Holmes EC, Kaidonis J, Lalueza-Fox C, la Rasilla de M, Rosas A, Semal P, Soltysiak A, Townsend G, Usai D, Wahl J, Huson DH, Dobney K, Cooper A. 2017. Neanderthal behaviour, diet, and disease inferred from ancient DNA in dental calculus. Nature.

23. Peabody MA, Van Rossum T, Lo R, Brinkman FSL. 2015. Evaluation of shotgun metagenomics sequence classification methods using in silico and in vitro simulated communities. BMC Bioinformatics 16:363.

24. Lindgreen S, Adair KL, Gardner PP. 2016. An evaluation of the accuracy and speed of metagenome analysis tools. Sci Rep 6:19233.

25. McIntyre ABR, Ounit R, Afshinnekoo E, Prill RJ, Hénaff E, Alexander N, Minot SS, Danko D, Foox J, Ahsanuddin S, Tighe S, Hasan NA, Subramanian P, Moffat K, Levy S, Lonardi S, Greenfield N, Colwell RR, Rosen GL, Mason CE. 2017. Comprehensive benchmarking and ensemble approaches for metagenomic classifiers 1–19.

26. Callahan BJ, McMurdie PJ, Rosen MJ, Han AW, Johnson AJA, Holmes SP. 2016. DADA2: High-resolution sample inference from Illumina amplicon data. Nat Methods 13:581–583.

27. Renaud G, Hanghøj K, Willerslev E, Orlando L. 2017. gargammel: a sequence simulator for ancient DNA. Bioinformatics 33:577–579.

28. Amir A, McDonald D, Navas-Molina JA, Kopylova E, Morton JT, Zech Xu Z, Kightley EP, Thompson LR, Hyde ER, Gonzalez A, Knight R. 2017. Deblur Rapidly Resolves Single-Nucleotide Community Sequence Patterns. mSystems 2:e00191–16.

29. Louvel G, Sarkissian Der C, Hanghøj K, Orlando L. 2016. metaBIT, an integrative and automated metagenomic pipeline for analysing microbial profiles from high-throughput sequencing shotgun data. Molecular Ecology Resources.

30. Huson DH, Auch AF, Qi J, Schuster SC. 2007. MEGAN analysis of metagenomic data. Genome Research 17:377–386.

31. Huson DH, Beier S, Flade I, Górska A, El-Hadidi M, Mitra S, Ruscheweyh H-J, Tappu R. 2016. MEGAN Community Edition - Interactive Exploration and Analysis of Large-Scale Microbiome Sequencing Data. PLoS Comput Biol 12:e1004957.

32. Nayfach S, Pollard KS. 2016. Toward Accurate and Quantitative Comparative Metagenomics. Cell 166:1103–1116.

33. Forster SC, Browne HP, Kumar N, Hunt M, Denise H, Mitchell A, Finn RD, Lawley TD. 2016. HPMCD: the database of human microbial communities from metagenomic datasets and microbial reference genomes. Nucleic Acids Res 44:D604–9.

34. Kennedy R, Lappin DF, Dixon PM, Buijs MJ, Zaura E, Crielaard W, O'Donnell L, Bennett D, Brandt BW, Riggio MP. 2016. The microbiome associated with equine periodontitis and oral health. Vet Res 47:49.

35. McDonald JE, Larsen N, Pennington A, Connolly J, Wallis C, Rooks DJ, Hall N, McCarthy AJ, Allison HE. 2016. Characterising the Canine Oral Microbiome by Direct Sequencing of Reverse-Transcribed rRNA Molecules. PLoS ONE 11:e0157046.

36. Sarkissian Der C, Pichereau V, Dupont C, Ilsøe PC, Perrigault M, Butler P, Chauvaud L, Eiríksson J, Scourse J, Paillard C, Orlando L. 2017. Ancient DNA analysis identifies marine mollusc shells as new metagenomic archives of the past. Molecular Ecology Resources 17:835–853.

37. Schnorr SL, Candela M, Rampelli S, Centanni M, Consolandi C, Basaglia G, Turroni S, Biagi E, Peano C, Severgnini M, Fiori J, Gotti R, De Bellis G, Luiselli D, Brigidi P, Mabulla A, Marlowe F, Henry AG, Crittenden AN. 2014. Gut microbiome of the Hadza hunter-gatherers. Nat Comms 5:3654.

38. Balvočiūtė M, Huson DH. 2017. SILVA, RDP, Greengenes, NCBI and OTT — how do these taxonomies compare? BMC Genomics 2014 15:1 18:114.

39. Green EJ, Speller CF. 2017. Novel Substrates as Sources of Ancient DNA: Prospects and Hurdles. Genes (Basel) 8:180.

40. Palmer SR, Miller JH, Abranches J, Zeng L, Lefébure T, Richards VP, Lemos JA, Stanhope MJ, Burne RA. 2013. Phenotypic heterogeneity of genomically-diverse isolates of Streptococcus mutans. PLoS ONE 8:e61358.

41. Truong DT, Tett A, Pasolli E, Huttenhower C, Segata N. 2017. Microbial strain-level population structure and genetic diversity from metagenomes. Genome Research 27:626–638.

42. Ahn T-H, Chai J, Pan C. 2015. Sigma: Strain-level inference of genomes from metagenomic analysis for biosurveillance. Bioinformatics 31:170–177.

43. Gonzalez A, Vázquez-Baeza Y, Pettengill JB, Ottesen A, McDonald D, Knight R. 2016. Avoiding Pandemic Fears in the Subway and Conquering the Platypus. mSystems 1:e00050–16.

44. Key FM, Posth C, Krause J, Herbig A, Bos KI. 2017. Mining Metagenomic Data Sets for Ancient DNA: Recommended Protocols for Authentication. Trends Genet.

45. Preus HR, Olsen I, Gjermo P. 1987. Bacteriophage infection—a possible mechanism for increased virulence of bacteria associated with rapidly destructive periodontitis. Acta Odontol Scand 45:49–54.

46. Preus HR, Olsen I, Namork E. 1987. The presence of phage-infected Actinobacillus actinomycetemcomitans in localized juvenile periodontitis patients. Journal of Clinical Periodontology 14:605–609.

47. Norman JM, Handley SA, Baldridge MT, Droit L, Liu CY, Keller BC, Kambal A, Monaco CL, Zhao G, Fleshner P, Stappenbeck TS, McGovern DPB, Keshavarzian A, Mutlu EA, Sauk J, Gevers D, Xavier RJ, Wang D, Parkes M, Virgin HW. 2015. Disease-specific alterations in the enteric virome in inflammatory bowel disease. Cell 160:447–460.

48. Kistler L, Ware R, Smith O, Collins M, Allaby RG. 2017. A new model for ancient DNA decay based on paleogenomic meta-analysis. bioRxiv 109140.

49. Lindgreen S. 2012. AdapterRemoval: easy cleaning of next-generation sequencing reads. BMC Research Notes 5:337.

50. Langmead B, Salzberg SL. 2012. Fast gapped-read alignment with Bowtie 2. Nat Methods 9:357–359.

51. DeSantis TZ, Hugenholtz P, Larsen N, Rojas M, Brodie EL, Keller K, Huber T, Dalevi D, Hu P, Andersen GL. 2006. Greengenes, a chimera-checked 16S rRNA gene database and workbench compatible with ARB. Applied and Environmental Microbiology 72:5069–5072.

52. Edgar RC. 2010. Search and clustering orders of magnitude faster than BLAST. Bioinformatics 26:2460–2461.

53. McMurdie PJ, Holmes S. 2014. Waste Not, Want Not: Why Rarefying Microbiome Data Is Inadmissible. PLoS Comput Biol 10:e1003531.

54. McMurdie PJ, Holmes S. 2013. phyloseq: an R package for reproducible interactive analysis and graphics of microbiome census data. PLoS ONE 8:e61217.

55. Lozupone C, Knight R. 2005. UniFrac: a new phylogenetic method for comparing microbial communities. Applied and Environmental Microbiology 71:8228–8235.

56. Lozupone CA, Hamady M, Kelley ST, Knight R. 2007. Quantitative and qualitative beta diversity measures lead to different insights into factors that structure microbial communities. Applied and Environmental Microbiology 73:1576–1585.

57. Foster ZSL, Sharpton TJ, Grünwald NJ. 2017. Metacoder: An R package for visualization and manipulation of community taxonomic diversity data. PLoS Comput Biol 13:e1005404.

